# A computational investigation on Rho-related GTP-binding protein RhoB through molecular modeling and molecular dynamics simulation study

**DOI:** 10.1101/2023.02.14.528547

**Authors:** Shamrat Kumar Paul, Chowdhury Lutfun Nahar Metu, Sunita Kumari Sutihar, Md. Saddam, Bristi Paul, Md. Lutful Kabir, Md Mostofa Uddin Helal

## Abstract

**Backgorund:** RhoB is a key member of the Rho family of isoprenylated small GTPases which modulate the cellular cytoskeletal organization. It has a crucial role in the neoplastic apoptotic mechanism after DNA damage. Due to the unavailability of 3D structure in the protein data bank database, in this study, we evaluated the structure of a protein, Rho-related GTP-binding protein RhoB.

**Results:** RhoB has a predicted pI of 5.10, indicating that it is acidic. The GMQE value was used to compute the target–template alignment, and 6hxu.1.A from Homo sapiens was chosen as the template structure, with the model construction task completed using swiss-model. The structural compactibility and stability were revealed after a 100ns molecular dynamics simulation using GROMACA employing the OPLS-AA force field. PCA analysis found residues that are relevant based on their fluctuation acitivity while their location is between 100-110 and 140-150.

**Conclusion:** This study will benefit future investigations addressing the association between gene mutation and abnormalities generated by protein Rho-related GTP-binding protein RhoB in apoptotic events by offering insight into the biophysical phenomenon of Rho-related GTP-binding protein RhoB inhibitors.

## 1. Introduction

Rho is one of the types of proteins weighing ∼21 kDa, which belongs to the Ras superfamily, characterized as “binary switches” in a diverse signalling pathway [1]. There are 3 subtypes of the Rho family including RhoA, RhoB and RhoC proteins that are 85% homogenous in their amino acid sequence identity, yet they each play distinctive biological roles in cell migration, and wound healing/ fibroelastosis or immune surveillance, cell division [2], [3]. Approximately, 20 signaling intracellular molecules are constituted by this family in humans and the dysregulation of their function has been linked to different human pathologies [4]–[8]. The cycle of RhoB continues through binding and non-binding of GTP/GDP which is similar to the other Rho GTPases. When GTP is in a bound state RhoB is considered an active state whereas a GDP-bound state is considered an inactive state [1], [9], [10].

RhoB can play both positive and negative regulatory roles in different physiology and pathophysiology. It remains at low steady-state levels in normal cell conditions and abnormal cells responding to several stimuli, including UV irradiation, growth factors, cytokines and phosphorylation during the cell cycle it can be rapidly and transiently upregulated [11], [12]. Because of their involvement in controlling cell migration and proliferation, Rho GTPases, especially RhoB, have been widely researched but least understood and still a mystery for their role in cancer progression [8], [13], [14]. RhoB regulates DNA damage responses, apoptosis, cell cycle progression, migration, and invasion, all of which contribute to cancer progression. During the progression of certain cancers, RhoB rates decrease. RhoB deficiency has been linked to increased cell proliferation, invasion, and metastasis [15].

RhoB is thought to be the most varied protein of the Rho subgroup, with multiple distinct post-translational modifications from RhoA and RhoC, possibly contributing to its bipolar role in cancer [8]. RhoB’s prominent involvement in vascular and endothelial cells likely leads to involvement in cancer vasculature, which seems to have consequences regarding angiogenesis [16]–[19]. The disparity in these findings has sparked controversy about whether RhoB promotes or inhibits tumour growth. These contradictory findings show that RhoB’s roles in cancer are highly contextual and cell-type specific. In the context of tumorigenesis vs tumour development and severity, numerous findings implied that RhoB performs two separate and opposing functions [3], [20]–[23].

Hence, the RhoB protein (mouse model) grabbed the attention of us to continue our findings. Unlike other organisms mice (*Mus musculus*) model has become a potent prop over many years in oncology research and our prior investigation of literature searching showed that homology modelling of the RhoB in mouse protein had not yet been investigated and well understood regarding the tumour and cancers. As it has mimicked human structure or physiology to know the RhoB protein expression using the mouse model is favourable. Considering the ethics of producing pharmaceuticals and medical treatments to treat diseases and illnesses, animals particularly mice are anticipated in scientific studies. Scientists may be able to dig up more prominent and similar medications, and treatments utilizing non-animal research approaches. If a new therapy appears to be promising, it is next tested in the human or human environment to see if it is both safe and effective. Human volunteers are asked to participate in a clinical trial if the outcomes of the animal studies are coming out positive. Animal studies are conducted initially to provide medical researchers with a better understanding of the benefits and drawbacks they might expect to see in people.

Since X-ray crystallography is considered as most trustable structure and its related study is time consuming, costly and tiresome which needed extensive effort and equipment; on other hand, in silico study gives some opportunities within a short period considering a few disadvantages. Therefore, to characterize the structural and functional properties of Rho-related GTP-binding protein RhoB and its encoded by the gene RhoB (Uniprot ID: P62746) consists of 196 amino acid long sequence to investigate its activity. Its subcellular location is the late endosome membrane; Cell membrane; Nucleus. For this purpose, we retrieved desired protein from Universal Protein Resource Knowledge Base (UniprotKB) [24] as a FASTA file format and due to the unavailability of 3-D structure in the Research Collaboratory for Structural Bioinformatics (RCSB) protein data bank (PDB) [25].

We analyzed the primary structure by ExPASy’s ProtParam tool [26] which showed its physicochemical features including molecular weight, GRAVY, theoretical pI, instability index (II) etc. For predicting the secondary structure elements SOPMA server was implied. Then the 3-D structure by homology modelling from SWISS-MODEL [27] relying on a central script-based software platform, ProMod3 [28] and this structure was qualified and analyzed from the Ramachandran plot by PROCHECK [29] program from SAVES v6.0, for identifying the compatibility of the atomic model (3D) with its amino acid sequences by VERIFY-3D. The ERRAT tool has been used to understand the non-bond interaction between each protein. Besides, we used ProSA-web [30] program for validating and checking the native protein folding energy of the model which was compared with the energy (potential mean force). Afterwards, to visualize the similarity of the template and model we used UCSF Chimera [31]. In addition, the last part of this computational investigation was accomplished by performing 100 ns (100,000 ps) Molecular Dynamics (MD) simulation in GROMACS platform with OPLS AA force field [32] and flexible water model spc216 that generated RMSD value, RMSF value and Radius of gyration (Rg) to identify the compactness of the model and reliability. Moreover, to visualize and identify the PC that grabs the attention in terms of analyzing the characteristics of the protein residue.

Furthermore, SASA gave information regarding the accessible surface area of the protein. Hierarchical clustering of the RMSD that gave the idea of the clusters that we assumed as PC and demonstrated in the time of 100,000ps through graphical presentation. To accomplished these tasks, we used the Bio3D function mktrj.pca.

## 2. Materials and Methods

### 2.1 Data Retrieval

From the UniProtKB database, we retrieved our desired protein Rho-related GTP-binding protein RhoB FASTA file. To understand at the amino acid level of the protein identified by the UniProtKB ID P62746 consist of 196 amino acid long sequence, and due to the unavailability of 3-D structure in the Research Collaboratory for Structural Bioinformatics (RCSB) protein data bank (PDB), this uncharacterised protein was taken up for predicting the 3-D structure by homology modelling approach as it is challenging and time-consuming to obtain experimental structures from X-ray crystallography or protein NMR method for every query protein. Proteins may fold in varied manners depending on energy conformation, steric factors, temperature, pH, concentration, etc. Hence, structure prediction is crucial to learning about the structural and functional characteristics of an uncharacterized protein.

### 2.2 Primary Sequence Analysis

The folding and intramolecular binding of the linear amino acid chain, which essentially defines the protein’s distinctive 3-D form, is driven by the primary structure of a protein from its amino acid sequence. The function of a protein is determined by its shape. The three-dimensional structure of a protein can be altered due to an alteration in the structure of the amino acid sequence, that protein becomes denatured and fails to fulfil its function as intended. ExPASy’s ProtParam tool; is a web-based server that computationally helps to determine the characteristic of a query protein. Physiochemical properties, include amino acid composition,hydropathicity, Molecular weight, number of atoms, Grade average of hydropathicity (GRAVY),Extinction coefficients, Theoretical pI, and Aliphatic index, which can give a clear definition and scenario of a protein. ExPASy’s ProtParam tool is reliable too which gives a clear understanding using its parameter about the physicochemical properties of Rho-related GTP-binding protein RhoB. Besides, the missing residue is also considered an important provision to know of a protein. PSIPRED 4.0 and DISOPRED3 [33] program.

### 2.3 Secondary Structure Analysis

The secondary structure evaluation of the query protein has been carried out with the help of the Self Optimized Prediction Method with Alignment (SOPMA) [34]. SOPMA is a tool that can predict the secondary structure of a query protein depending on the primary sequence of a protein, using Window width: 17, Similarity thresholds: 8, Number of states: 4; parameters.According to this methodology, a short homologous sequence of amino acids tends to form a similar secondary structure which shows whether it lies in a helix, strand or coil.

### 2.4 Homology Modeling

Homology modelling harnesses a sophisticated method to elucidate the experimental data with high accuracy in the computational biology, pharmacological and biomedical research fields [35]. Since the experimentally solved structure of Rho-related GTP-binding protein RhoB is not available in (RCSB) PDB, to elucidate in-silico biophysical characteristics, the 3-D structure has been predicted by using homology modelling became utmost. The structure prediction of our protein of interest was accomplished by the SWISS-MODEL [36] program.

### 2.5 Validation and Quality Check

The quality of the model was validated to examine its credibility and consistency by a few tools. PROCHECK analysis quantifies the available zone of residues presented by checking the Ramachandran plot [37]. Along with, the VERIFY-3D [38] program to identify the compatibility of the atomic model (3D) with its amino acid sequence (1D). All of that analysis was brought out based on structure Analysis and Verification Server (SAVES) v6.0. The overall quality factor of the query protein was being checked with the non-bond interaction between each protein, ERRAT [39] tools had been used, and subsequently, PorSA-web [40] program was engrossed in validating and refining to check the native protein folding energy of the model which compared with the energy (potential mean force) derived from a long set of protein structures. For checking the superimposition by UCSF Chimera [31] applied to visualize the similarity of template and model.

### 2.6 Molecular Dynamics Simulation

All-atom molecular dynamics (MD) simulations are a powerful tool for investigating biomolecular motion on the pico- to nanosecond time scale. While visualizing an MD trajectory provides qualitative insights into the system, quantitative data that describes or supports these insights are often difficult to obtain [41]. So, It is necessary to know the nutshell of the stability, endurance and dynamics of the modelled protein MD simulations were led using OPLS-AA force field and flexible water model in GROMACS 2021.2 [42] package continued in Linux OS (Ubuntu20.04 distribution). At that time all atoms of 6hxu.1. A protein was enclosed by a water box called genion of spc216 water model. When it was being solvated, the system was neutralized with 4Na+ ions. That solvated system then minimized the energy through 5000 steps of the steepest invasion to eradicate the van der Waals contacts initiating a favourable structure for MD simulation. Using the LINCS algorithm [43], all bonds were constrained. Furthermore, the electrostatic bond interactions were elucidated by the particle mesh Ewald (PME) algorithm [44].

### 2.7 Essential Dynamics Analysis

Principal component analysis (PCA) was applied to study conformational flexibility by employing a basic statistical technique called PCA to understand the collective motions of Rho-B. PCA is the process of computing the principal components and using them to change the basis of data,often simply using the first few and disregarding the rest. [41]. PCA is a method for detecting and identifying patterns in high-dimensional data and displaying them graphically to highlight their similarities and differences. The Bio3D function mktrj.pca was used to do the PC analysis [45]. Visual inspection of interpolated atomic displacement trajectories along with the first three PCs. Least-squares fitting to the first frame reduced the trajectory’s total translational and rotational motions. Using Cartesian coordinates of C atoms, a 3 N 3 N covariance matrix was created. The covariance matrix was diagonalized, resulting in 3 N eigenvectors, each with its eigenvalue. The covariance matrix is specified in terms of deviations from the averaged coordinates of the trajectory, and the RMSF plot can be shown using eigenvectors. When computing principal component scores, it is preferable to decrease their number to an independent set, whereas eigenvectors are the weights in a linear transformation. The amount of variance explained by each primary component or factor is expressed in eigenvalues. The number of components recovered is equal to the number of observed variables in the analysis, and each Principal Component has the same amount of variance. The first principal component identified accounts for most of the variance in the data. The second component identified accounts for the second-largest amount of variance in the data and is uncorrelated with the first principal component and so on [41] For PC analysis, we employed a 10ns trajectory with 1000 frames. We have followed the PCA analysis protocol described on Paul et al. 2022 for this approach [46]. To reveal concerted motions, the trajectory was projected onto a certain eigenvector. K-means and hierarchical clustering methods were used to cluster the trajectory in the PC space. By reducing the mean squared distance between each observation and its nearest cluster centre, k-means splits the observations into k clusters. The trajectory can be projected on eigenvectors to give the principal components pi(t) [45], [47]:

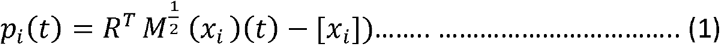

### 2.8 System Details

The Primary structure from ExPASy’s ProtParam tool, Disopred plot from PSIPRED 4.0; DISOPRED3 program, Secondary structure from SOPMA, Clustal Omega [48] for Multiple sequence alignment 3-D structure from SWISS-MODEL through ProMod3 3.2.0 web-server and structure validation analysis using PROCHECK v.3.5.4 [29], [49], Verify 3-D, ERRAT and PorSA-web tool from SAVES v6.0 [50], were carried out in online. Superimpose structure visualization in UCSF Chimera was been performed on Hp ProBook 645 G4; AMD RAYZEN 7pro 2700U. Molecular dynamics simulation was carried out in Science Outreach Server’s GROMACS program via SSH login mode on the local windows subsystem for the linux command line interface. The Bio3D function mktrj.pca for calculating PC, SASA and cross-validation to earn the knowledge to earn the total surface area, hydrophobicity index and residue x residue index [45], [47].

## 3. Result and Discussion

### 3.1 Primary Sequence Analysis

The amino acid sequence of Rho-related GTP-binding protein RhoB was retrieved from the UniprotKB database in FASTA format which was exploited for computing physicochemical properties from ExPASy ‘s ProtParam tool using its parameter and the subsequent retrieval of Molecular weight, theoretical, Isoelectric Point (pI), GRAVY, and aliphatic Index of query protein was were calculated that are tabulated in **Table 1**. The calculated molecular weight of 22123.39 Da for the query protein. The stability of a protein is measured in a test tube depending on its instability index [51]. The instability index of a protein and its stability isn’t complimentary, if a protein shows its instability index of more than 40, it can’t be able to stable in an in vivo method of more than 5hrs but a protein with less than 40 shows its stability approximately 16 hrs [52]. **Table 1** showed that the instability index of our query protein is 46.35 [53] which classified it as unstable but merging with the threshold line which indicates the probability of a significant level of stability. The theoretical pI of this query protein is 5.10 which indicates its acidic nature whereas the pI of a protein less than 7 is supposed to be acid and shows its acidic nature [51]. A statistical analysis experimented upon the proteins of thermophilic bacteria that showed a higher aliphatic index than that of ordinary proteins. The aliphatic index was defined as the relative volume of a protein occupied by aliphatic side chains (alanine, valine, isoleucine,and leucine). This index may be regarded as a positive factor for the increase of the thermostability of globular proteins. An aliphatic index of 87.96 demonstrated that protein had a positive factor in the increase of thermostability [54]. The enumerated GRAVY was -0.60 as shown in **Table 1** conveying its hydrophobicity nature shows the possibility of its better interaction with water [55], [56]. From **Figure 1A**, among all the amino acids, Valine (Val)residue’s participation was higher around 25% than Aspartic acid (Asp) and Glutamic acid (Glu)contributed more than 15% respectively. The configuration of a protein is generally derived from X-ray diffraction data. If the experimentalists are unable to solve the structure of that area by using X-ray diffraction data, those regions will mark as missing residues in the PDB file. The long-chain amino acid sequence contains more than thirty amino acids, around 1/5th of a typical eukaryote proteome, and that 1/3rd of eukaryotic proteins comprise the nearby disordered region and they comprehend a few important regulatory processes which were suggested by conservative estimates of the disorder frequencies in complete genomes [57].

**Table 1.** Physicochemical properties of RhoB protein computed using EXPASY’s Protparam tool.

| Properties | Number |
| --- | --- |
| Number of Amino Acids | 196 |
| Molecular weight | 22123.39 |
| Theoretical pI | 5.10 |
| Total Number of atoms | 3103 |
| Aliphatic index | 87.96 |
| Total number of negatively charged residues (Asp + Glu) | 32 |
| Total number of positively charged residues (Arg + Lys) | 26 |
| Extinction Coefficient; Assuming all pairs of Cys residues form cystines ( M-1 Cm-1 ) | 21930 |
| Extinction Coefficient; Assuming all Cys residues are reduced (M-1 Cm-1 ) | 21430 |
| Instability Index (II) | 46.35 |
| Grade average of hydropathicity (GRAVY) | -0.60 |
| Half-life; mammalian reticulocytes, in vitro (hours) | 30 |

**Figure 1:**
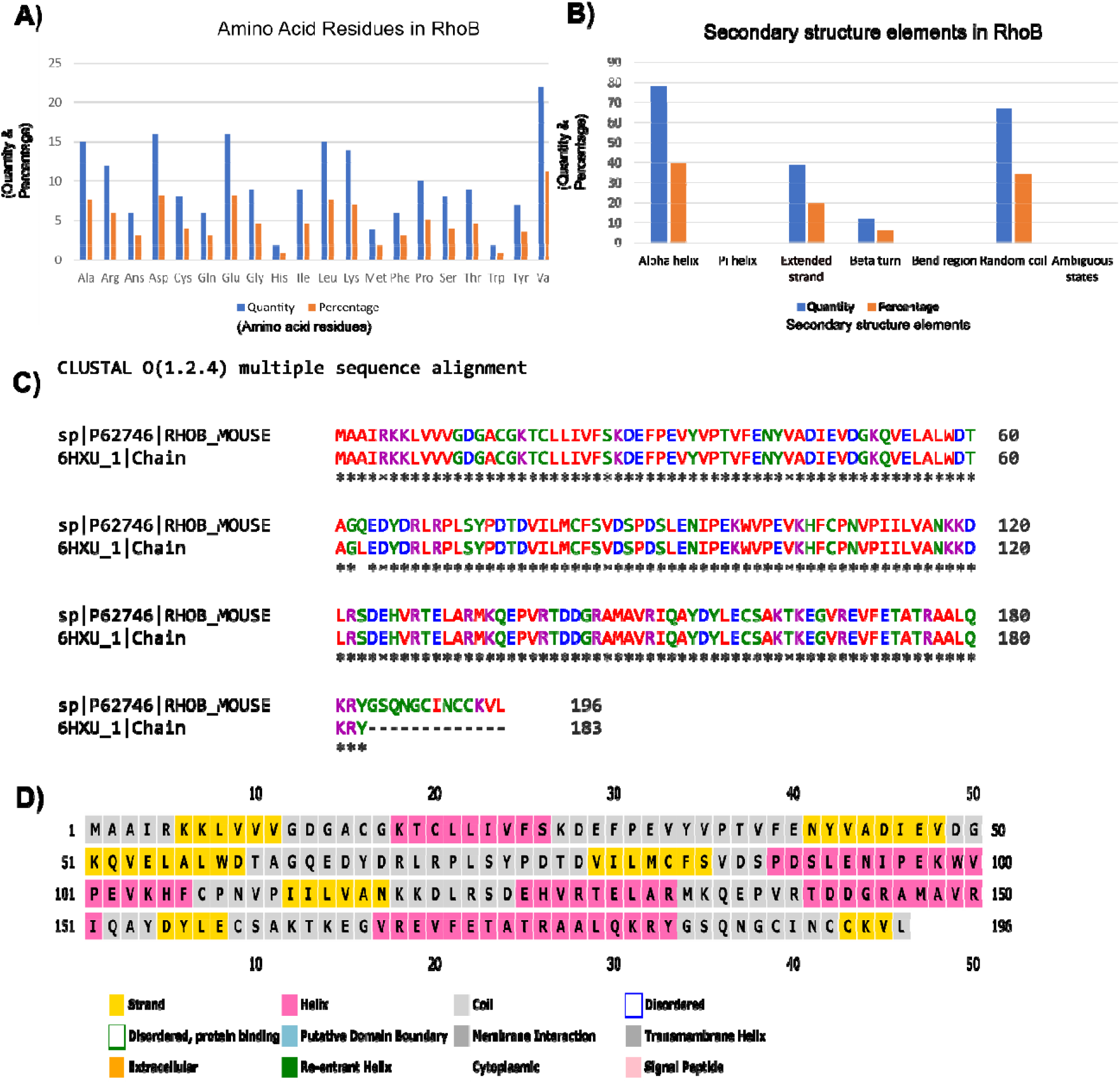
Primary sequence, secondary structure elements, disorder residues and multiple sequence analysis. **(A)** Among all the amino acids, Valine (Val) residue’s participation shows the higher value that reaches around 25% consequently Aspartic acid (Asp) and Glutamic acid(Glu) shares more than 15% of residues respectively. **(B)** Contribution of secondary structures are showed in percentages, 34.18% of residues as a random coil(cc) compare to 78% to Alpha helix, extended strand (Ee) 19.9% and β turn (Tt) is 6.12% which estimated it as α helix dominated protein. **(C)** Multiple sequence alignment (MSA) analysis performed between query protein sequence and template protein sequence, 6HXU.1. A by Clustal Omega tool which possesses the 100% identity between two sequences. **(D)** Disordered residues of the protein has been depicted here while white colour box represents disordered residue and there are no existing disordered amino acid residues according to DISOPRED prediction.

### 3.2 Secondary Structure Analysis and Model Building

The secondary structure of query protein was predicted from SOPMA. The success rate in the prediction of the secondary structure of proteins of 69.5% accuracy of amino acids for a three dimensional-state. Description of the secondary structure contains α-helix, β-sheet and coil in a whole protein [58]. SOPMA prediction uses the parameters of window width 17, the number of states 4 and similarity threshold 8 that were used for the secondary structure prediction. According to **Figure 1B**, secondary structure prediction by SOPMA shows each amino acid’s contribution in each position to the secondary structure 34.18% of residues as a random coil(cc) compare to 78% to Alpha helix, extended strand (Ee) 19.9% and β turn (Tt) is 6.12% which estimated it as α helix dominated protein.

Multiple sequence analysis (MSA) can provide inferred and phylogenetic analysis which can be elucidated to assess the sequences’ shared evolutionary origins. The multiple sequence alignment can predict the sequences are homologous, they descend from a common ancestor. The algorithms will try to align homologous positions or regions with the same structure or function. Clustal Omega [59] uses seeded guide trees and Hidden Markov Model (HMM) profile-profile techniques to generate alignments. Furthermore, The retrieval of a homologous sequence to construct an alignment via Basic Local Alignment Tool (BLAST) Programs [60], [61]. The sequence alignment **Figure 1C** target-template; with 100% identity; indicated the amino acid residues from target and template sequences shared identical amino acid regions and it imposed perfectly with our query protein that will show the physicochemical properties as like the template protein. Secondary structure information and feature data helped to make an accurate alignment and it served as the model of the protein domain. In **Figure 1D** white colour represents disordered residue and there are no existing disordered amino acid residues according to DISOPRED prediction.

Homology modelling is one of the premise processes to predict the 3-D structure of an unknown protein also known as comparative modelling of protein. A central script-based software platform, ProMod3 [62] has been used to generate the model through the SWISS-MODEL web server [36]. It was performed under a few steps, firstly template structure selection before submission of query protein sequence in FASTA format. The predicted protein was selected depending on the GMQE score, and identification. From the SWISS-MODEL template library, we got more than 30 templates found by the HHblits method [63] which is profiled by Hidden Markov models (HMMs) the fastest sequence search tool [64] among them 5 most favourable templates is 2fv8.1.A, 6sge.1.A, 2fv8.1.A, 6sge.1.A, 6hxu.1.A (**Table 2**). Considering the Global Model Quality Estimation (GMQE) a method of quality estimation that calculates both the target–template alignment and the template structure ranging from 0 to 1 indicating the predicted accuracy of a model designed with that alignment and template, as well as the target 6hxu.1.A, coverage score, 0.93 with 99.45% identity considering GMQE value of 0.85, possesses reasonably higher score among those 5 templates. We put forward our work with 6hxu.1. A protein template from *Homo sapiens*. Furthermore, we proceeded with our work with a favourable GMQE score, identity and method (X-ray, 1.19A) as an ideal protein prediction template for our purpose. Modelling a protein based on predicted template structure relies on the thread method close to the 3D structure of a stranded protein where a complete structure of a protein atom, as well as folds protein, is also being collected for a possible set of templates [65]. To Visualize the structure of qualified protein, we built the model depicted in **Figure 2A** from SWISS-MODEL.

**Table 2.** Template selection for modeling RhOB (found by HHblits) through expasy’s swiss-model tool.

| Description | Organism | PDB ID | GMQE | Coverage | Identity(%) | Method |
| --- | --- | --- | --- | --- | --- | --- |
| The crystal structure of RhOB in the GDP-bound state | Homo sapiens | 2fv8.1.A | 0.81 | 0.94 | 100 | X-ray<br>1.90 Å |
| Crystal structure of Human RHOB-GTP in complex with nanobody B6 | Homo sapiens | 6sge.1.A | 0.84 | 0.93 | 99.45 | X-ray<br>1.50 Å |
| The crystal structure of RhOB in the GDP-bound state | Homo sapiens | 2fv8.1.A | 0.81 | 0.93 | 100 | X-ray,<br>1.90 Å |
| Crystal structure of Human RHOB-GTP in complex with nanobody B6 | Homo sapiens | 6sge.1.A | 0.84 | 0.93 | 99.45 | X-ray,<br>1.50 Å |
| Crystal structure of Human RHOB Q63L in complex with GTP | Homo sapiens | 6hxu.1.A | 0.85 | 0.93 | 99.45 | X-ray,<br>1.19 Å |

**Figure 2.**
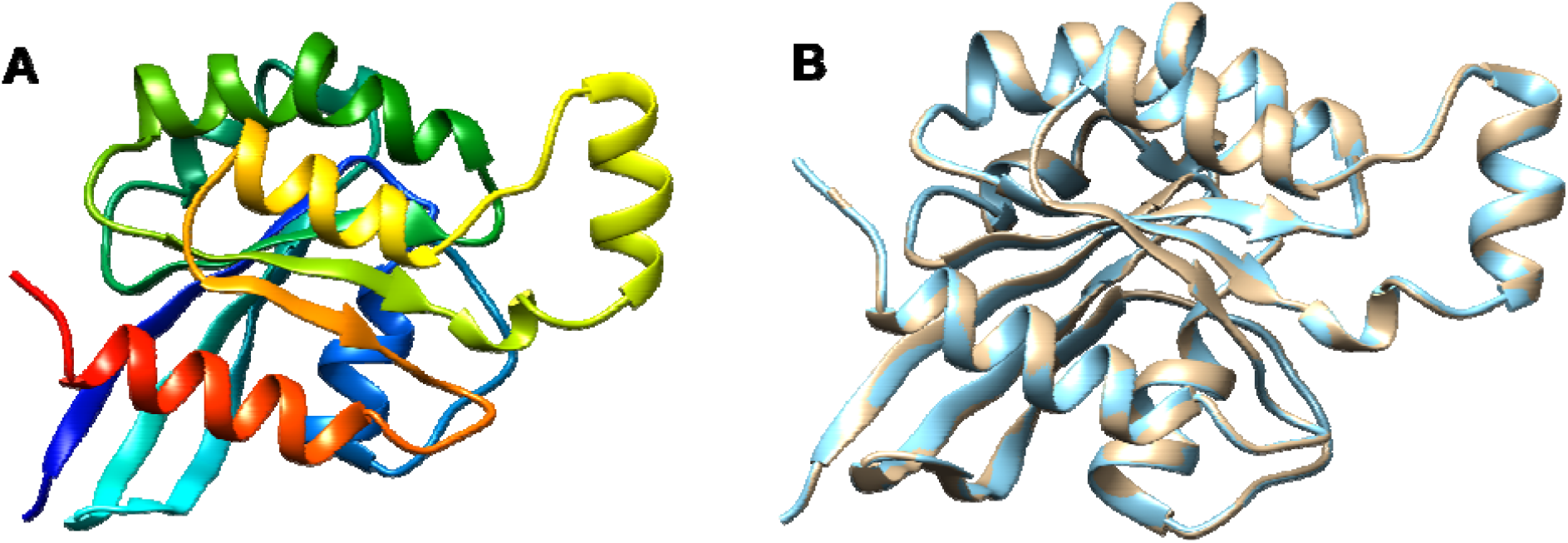
Predicte Structure of modeled protein and the superimposed structure with template, 6hxu.1. **A**. (A) The depicted modeled structure representing from N-terminal to C-terminal where red color dignifies Helix, green color Coil and Strands by yellow color. (B) Superimposed structure between modeled protein of RhoB and the template protein 6hxua.1.A, showed by golden color and Model by light blue color. This structure indicates that the selected template and predicted model superimposed perfectly. Visualization of this structure performed by UCSF Chimera program.

For moving forward, identify the template and the model whether they were superimposed or not (**Figure 2B**). For this purpose, UCSF Chimera gave better visualization for observing the superimposed structure for the query protein. Template of the protein (golden colour) and model (light blue colour) both that were required from the SWISS-MODEL, entirely showed superimpose structure in UCSF Chimera that was seemingly favourable. The sequence of amino acid residues comprising a binding pocket or cavity site defines its physicochemical properties, which along with its form and position in a protein, describe its functionality. Protein mobility allows for the opening, closure, and adaptation of binding pockets to control binding processes and basic protein function [66]–[68].

### 3.3 Validation and Quality Check

PROCHECK; Saves v6.0, an online server tool that Checked the stereochemical quality of a protein structure by analyzing residue-by-residue **Figure 3 (A and B)** gives overall structure geometry, the same resolution and also highlighted the regions that may need further investigation. The number of residues in allowed and generously allowed regions [A, B, L] was 149; 92.0% (model) and 149; 91.4% (template) respectively and none of the residues were present in the disallowed region of the plot (**Table 3**). No greater than 20%, a good quality model would be expected to have over 90% in the most favoured regions. So, we assumed our selected template can probably give a well-structured prediction model. The predicted 3D structure was validated on the studied Protein 6hxu.1.A using PROCHECK program [69], which estimates phi/psi angles, constructed Ramachandran plots. The quality of our structure again verified by the ERRAT score was 98.802 **(Figure 3C)** for the model and the template shows an overall quality factor of 94.737 **(Figure 3D)**; indicating an acceptable environment for protein. Moreover, after building the model from the selected template we got a few parameters such as Z-score known as a standard score that indicates how far a data point deviates from the baseline. A protein’s Z-score is defined as the energy separation between the native fold and the average of an ensemble of misfolds in units of the ensemble’s standard deviation [70], [71]. ProSA-web program that profiled the energy of the template and model and the Z-score value,is a parameter to measure the quality of a model as it quantifies the total energy of the structure. This Z-score value was obtained from by ProSA-web program that measured the interaction energy of each residue and applied a distance-based pair potential. The score of **-6.27** for the template and -6.54 for the model **(Figure 3E, 3F)** was analysed by the ProSA-web program where the negative ProSA energy reflects the reliability of the model [51]. Using the VERIFY-3D tool (**Figure 4)** showed that the model passed the requirements and it demonstrated at most of the amino acids had positive values and resides above the value of 0.5 both for the model **Figure 4A** and template protein **Figure 4B** indicating endurance and reliability.

**Table 3.** Ramachandran Plot Statistics of Model and Template Structures.

|  | Plot Statistics | Model | Template |
| --- | --- | --- | --- |
| Region | Residues in most favoured regions [A,B,L] | 149 (92.0%) | 149 (91.4%) |
|  | Residues in additional allowed regions [a,b,l,p] | 13 (8.0%) | 14 (8.6%) |
|  | Residues in generously allowed regions [~a,~b,~l,~p] | 0 (0%) | 0 (0%) |
|  | Residues in disallowed regions | 0 (0%) | 0 (0%) |
|  | Total | 0 (0%) | 100% |
| Residues | Number of non-glycine and non-proline residues | 162 | 163 |
|  | Number of end-residues (excl. Gly and Pro) | 2 | 2 |
|  | Number of glycine residues (shown as triangles) | 7 | 7 |
|  | Number of proline residues | 10 | 10 |
|  | Total number of residues | 181 | 182 |

**Figure 3:**
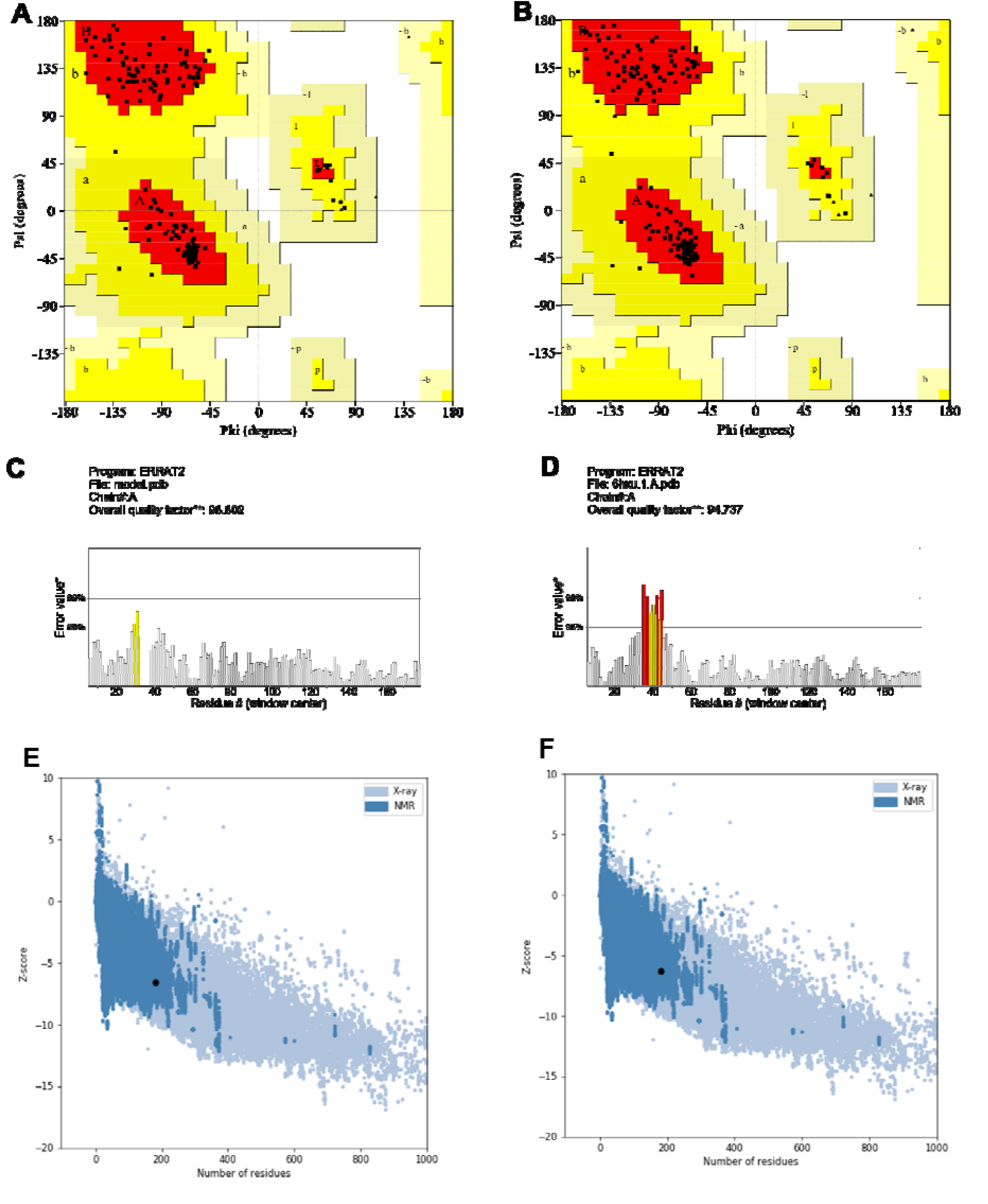
Ramachandran plot, ERRAT value and Z-Score. Structure validation through Ramachandran plot analysing (PROCHECK; Saves 6.0), **(A)** dignifies the modeled protein Ramachandran plot plot and **(B)** depicted the template, 6hxua.1.A. The number of residues in allowed and generously allowed region [A, B, L] 149; 92.0% (model) and 149; 91.4% (template) respectively. The quality of our structure again verified by ERRAT score was 98.802 **(C)** for the model from and the template shows an overall quality factor of 94.737 **(D);** Based on the condition it can say that our template and model has less error or rejection value. Z score of the Template and Model. The score -6.54 **(E)** for the template -6.27 **(F)** analysed by the ProSA-web program where the negative ProSA energy reflects the reliability of the model.

**Figure 4.**
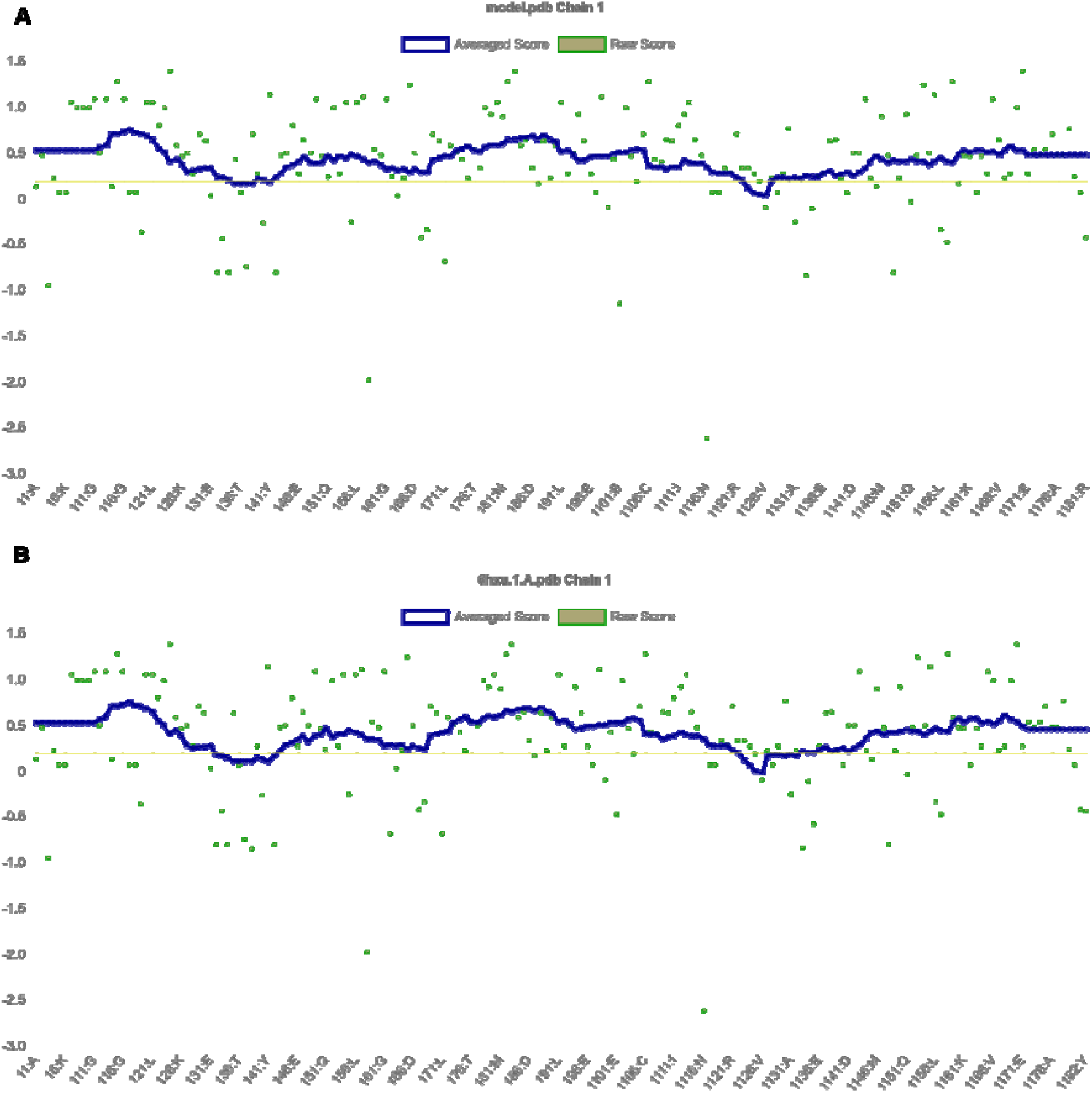
Verify 3D tool fix the compatibility of an atomic model in 3D structure level with its own amino acid sequence in primary form. Using the VERIFY-3D tool resulting graph shows that the model passed the requirements and it demonstrated at most of the amino acids had positive value and resides above value 0.5 both for the template and modelled protein indicating endurance and reliability.

### 3.4 Molecular Dynamics Simulation

The structure was further validated against energy minimization by MD simulation using GROMACS on Ubuntu-2021.2 applying an all-atom optimized potential for liquid systems (OPLS-AA) force field [72]. MD simulation trajectories were analysed to obtain GROMACS energies, root square deviation (RMSD), root-mean-square fluctuation (RMSF), and the radius of gyration (Rg); values are depicted in **Figure 5**. Protein’s potential energy was -7e10^6^ Kj mol^-1^, but it reduced during the simulation to nearly -1×10^6^ depicted in **Figure 5A**. Electrostatic cut-off and Vander Waals cut off value was 1.0 and 1.0 respectively. In the equilibration state NVT and NPT; NVT started to run for 50000 steps and for 100.0 ps to restrain the position of the protein and based on the parameter of leap-frog integrator, where the initial temperature was 298.948K. NVT (N expressed as a constant number, V volume and energy (E); and it is the sum of kinetic (KE) and potential energy (PE) which is conserved and T and P are unregulated equilibrations generated temperature **(Figure 5B)** showed the higher and lower temperature peak in that particular graph and the highest value was around 303K. In NPT (constant number N; Pressure P and Temperature T) equilibrium process it conducted the pressure until it came to the proper density. In **Figure 5C** the x-axis is considered as time in ps and y-axis as pressure showed the highest and lowest value was nearly 300 bar and -220 bar respectively whereas **Figure 5D** shows the density: highest peak along y-axis was around 1023 Kg m^-3^ and the lowest value was around 1010 Kg m^-3^.

**Figure 5.**
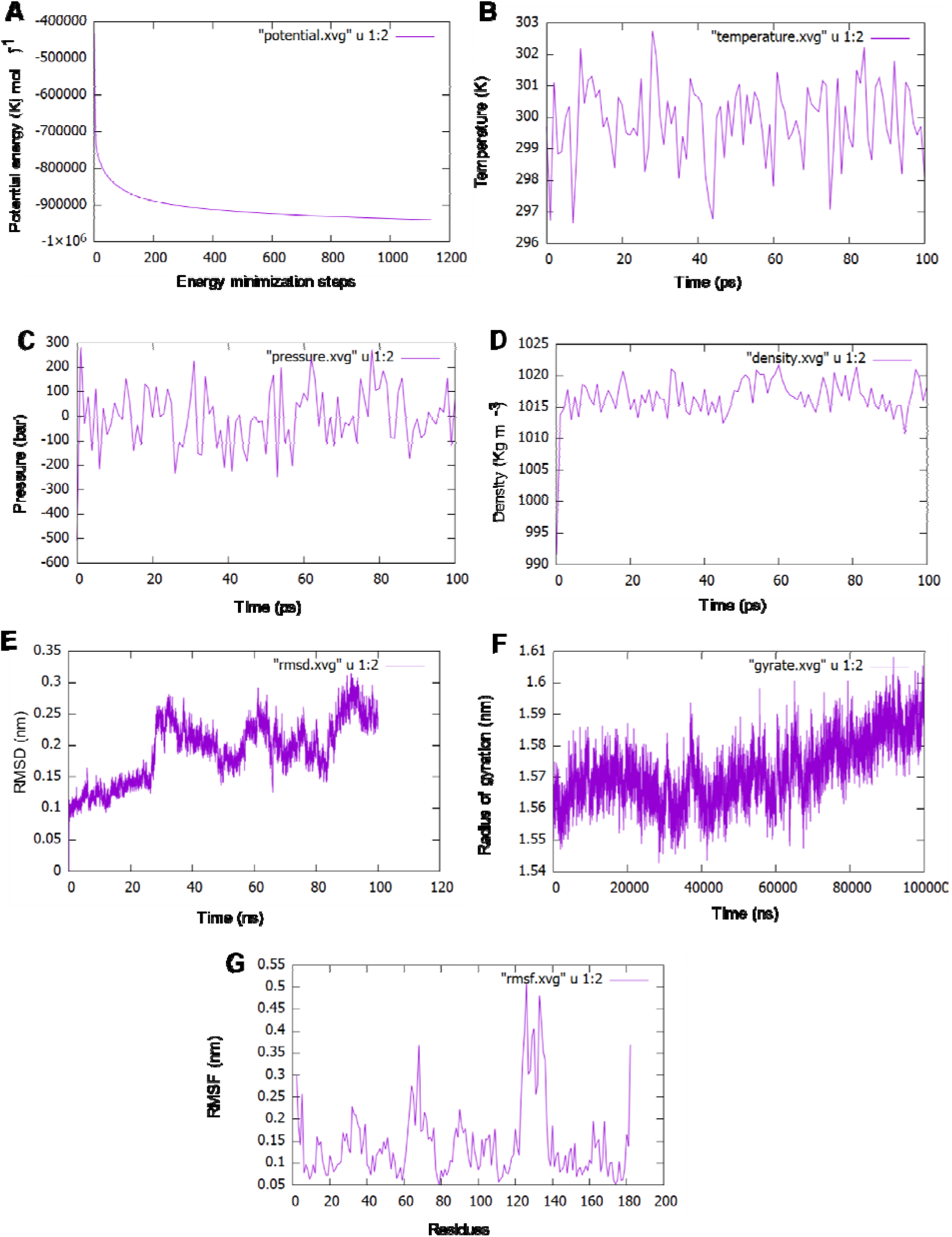
100 ns molecular dynamics (MD) simulation of RhoB protein. **(A)** Protein’s potential energy was -7e10^6^ Kj mol-^1^, but it reduced during the simulation nearly -1×10^6^. **(B)** The equilibrations generated temperature showed the higher and lower temperature peak in that particular graph and the highest value was around 303K. **(C)** The pressure graph showed the highest and lowest value of pressure fluctuation in the entire simulation which was nearly 300 bar and -220 bar respectively. **(D)** the density trajectory shows highest peak along was around 1023 Kg m^-3^ and the lowest value was around 1010 Kg m^-3^. **(E)** A molecular dynamics simulation was running for performing 100 nanoseconds (100,000ps) where the lowest and highest RMSD value is 0.1 nm and nearly 0.36 nm accordingly. **(F)** The Radius of gyration (Rg) graph depicts that atoms were fluctuated highest nearly 1.65 nm at around 95ps and the lowest fluctuated value was almost 1.54 nm around 30ps. **(G)** The RMS Fluctuations, of each atom echo, generated shows that residues range between 120 and 140 (amino acid sequences: DLRSDEHVRTELARMKQEPVR) fluctuated mostly towards 0.5 nm. The RMS fluctuation figure reveals also the C-alpha backbone variance during the simulation process.

A molecular dynamics simulation was running for performing 100 nanoseconds (100,000ps). The lowest and highest RMSD value (**Figure 5E)** is 0.1 nm and nearly 0.36 nm accordingly. During the period of 0 to 30 ns, the trajectory raises steadily and it fluctuates in an orderly for the rest of the period till it reaches. The Radius of gyration (Rg) graph shown in **Figure 5F** depicts those atoms were fluctuated highest at nearly 1.65 nm at around 95ps and the lowest fluctuated value was almost 1.54 nm at around 30ps. That showed atoms of our query protein did not fluctuate much and evaluated that this structure was in a compact condition in contrast the RMSD value represented the reliability of the structure. The RMS Fluctuations, of each atom echo, generated showed in **Figure 5G** showed that residues range between 120 and 140 (amino acid sequences: DLRSDEHVRTELARMKQEPVR) fluctuated mostly towards 0.5 nm. The RMS fluctuation figure reveals also the C-alpha backbone variance during the simulation process. The tolerable fluctuations in the backbone that proven the reliability of protein models [73].

### 3.5 Principal Component Analysis

A molecular dynamics simulation was run for 100 ns before actually principal component analysis (PCA). Components that account for the most variance are kept, while those that account for a small amount of variance are removed. In molecular dynamics (MD) simulations, PCA and clustering are used to reveal substantial conformational changes. [41]. Only a small part of these PCs describes a great majority of the whole atomic movement [47], [74] and these motions are often essential for protein function [75].

As a result, it seemed sensible to use the PCA method to reduce the phase space of proteins for long-term molecular dynamics [76]. For this purpose, PCA used here to identify a small number of important modes and then project the equations of motion onto the resulting low-dimensional vector space. Due to this crystal structure of RhoB’s and immersion in water, the model required a significant amount of equilibration time. The system’s RMSD and radius of gyration; Rg were monitored for 100ps to establish equilibration.

In addition, both for the entire 10,000 frames in the 100ns time frame as well as for the last 1000 frames was considered to capture principal components. The resulting data matrix was subjected to principal component analysis. Initially, we considered whole 10,000 frames of entire hundred nanoseconds of the alpha-carbon chain (181 atoms, 181 residues) to calculate the maximum pairwise RMSD for clustering **(Figure 6A)** and it was calculated as 4.905646 nm. RMSD hierarchical clustering demonstrates the hierarchical distribution of the total clusters by colours code ranging from red, blue, sky blue, purple, black, green and yellow. In this hierarchical clustering, we can observe the yellow clustering which appeared also in the Pairwise PC analysis and the Cartesian coordinate indicates the location of the clustering residues in the last stage of the entire 100, 000ps of simulation. Therefore, cartesian coordinate PCA analysis **(Figure 6B)** reflects some extent the dominant overall motion rather than the much smaller internal motion of the modelled RhoB protein [77]. **Figure 6B** describes the cartesian movements of PC that are manipulated as the overall motion of the protein. Whereas, pairwise distance PCA (**Figure 6C**) in the distance pairwise PC analysis, we were able to predict the position of atoms over time. In this 2D graph a colour range was included and we observed in the starting they were scattered between the 1^st^ and 4^th^ coordinate but in the terminate state, they started to appear in the 2^nd^ and 3^rd^ coordinate which was marked by yellow colour.

**Figure 6.**
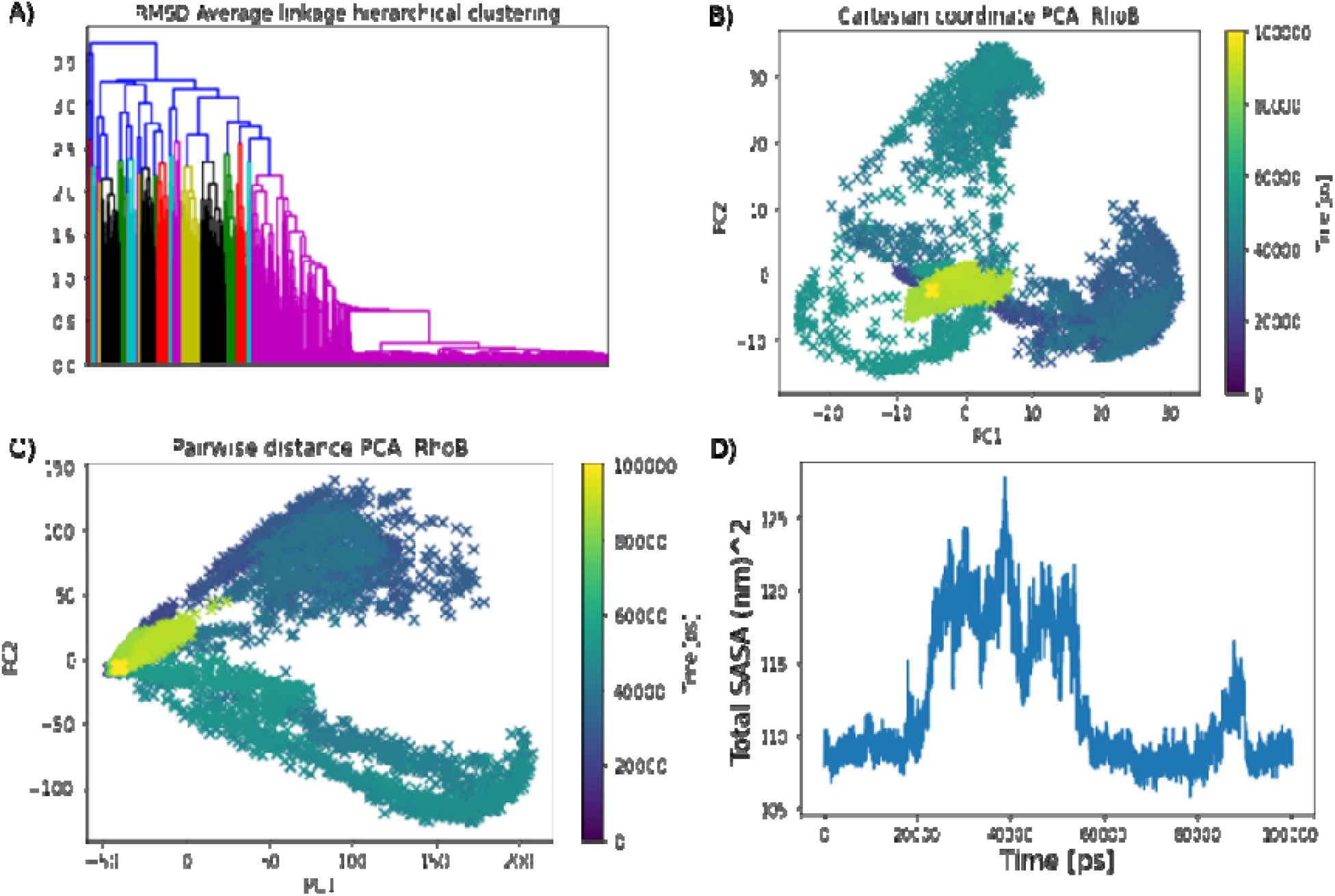
Principal component analsysis for entire 10,00 frames. **(A)** RMSD Average linkage hierarchical clustering depicts the movements of atoms with respect to time from starting to termination state in a whole for 100ns. (**B**) The cartesian coordinate PCA analysis reflects the dominant overall motion to some extent rather than the much smaller internal motion of the modelled RhoB protein. (**C**) The prediction of atom positions over time using pairwise distance PCA. (**D**) The fluctuation of Solvent Accessible Surface Area (SASA) during the simulation at 300 K for 100ns.

To further understand Rho-B correlated dynamics, we considered, Solvent accessibility (SASA) is a key feature of proteins for determining their folding and stability [78]. To compute the solvent-accessible surface area of each atom or residue in each simulation frame. In our analytical part, SASA analysis revealed the solvent-exposed area, which may decrease the solubility of the protein and can modulate protein-protein interactions [3], [79]–[81]. Prediction of protein solubility is gaining importance with the growing use of protein molecules as therapeutics, and ongoing requirements for high-level expression [75]. **In Figure 6D** we can see the fluctuation of total atoms; 181 residues plotted against 10,000 frames for 100ns (100,000ps). The number of points representing the surface of each atom, higher values lead to more accuracy. Assuming that the points are evenly distributed, the number of points is directly proportional to the accessible surface area. From the GetArea web tool [77] we calculated the SASA and it showed a total surface 9514.27 **(Table 4)**.

**Table 4.** Calculated properties from Solvent Accessible Surface Area (SASA)

| Parameters | Numbers |
| --- | --- |
| POLAR area/energy | 3756.87 |
| APOLAR area/energy | 5757.40 |
| UNKNOWN area/energy | 0.00 |
| Number of surface atoms | 797 |
| Number of buried atoms | 653 |
| Total area/energy | 9514.27 |

However, PC analysis for the whole simulation time frame can miss relevant motions since major collective dihedral transitions do not usually correspond to major transitions in Cartesian space. Therefore, principal component (PC) analysis was performed using Cα Position covariance while we considered terminal 1000 frames and analyzed principal components using the **Bio3D package** of R [45], [47] to look for further insight into the last 1000 frames. The atomic coordinates of the Cα atoms (181 total atoms) were used in the analysis to reduce statistical noise and to avoid artificial apparent correlations between slow side-chain fluctuations and backbone motions. A PCA in dihedral angle space is based on internal coordinates which naturally provide a correct separation of internal and overall motion.

**Figure 7A** elucidated through an average fluctuation graph of a total of 181 components and a principal component. In this graph along the x-axis the position of residues and along the y-axis the PC was subjected and the rate of fluctuation of PC was higher in the position ranging from 100-150 (Å) with blue colour and the average fluctuation rate is in between (0.10). So, residues that hold the position between 100-110 and 140-150 are phenylalanine, glutamate, valine, lysine, proline, histidine, cysteine, asparagine, arginine threonine, aspartic acid, glycine, alanine, methionine, alanine, valine and arginine that we got from our multiple sequence alignment. These residues take into account. From the coloured graph (**Figure 7B**) we observed that clusters are occupied within all 4 coordinates and in some places, they are highly dense and on the other side they are not much dense as previous. This dense place is considered as the cluster with the same features or influence of subspace. Variables contributing similar information are grouped, that is, they are correlated. When the positively correlated numerical value of one variable increases or decreases, the numerical value of the other variable tends to change in the same way. When variables are negatively (“inversely”) correlated, they are positioned on opposite sides of the plot origin, in diagonally opposed quadrants [82]. Furthermore, the distance to the origin also conveys information. The further away from the plot origin a variable lies, the stronger the impact that variable has on the model. In **Figure 7C**, it’s shown the contributions of the first PC in a structural form with blue and red colours through a ribbon structure.

**Figure 7.**
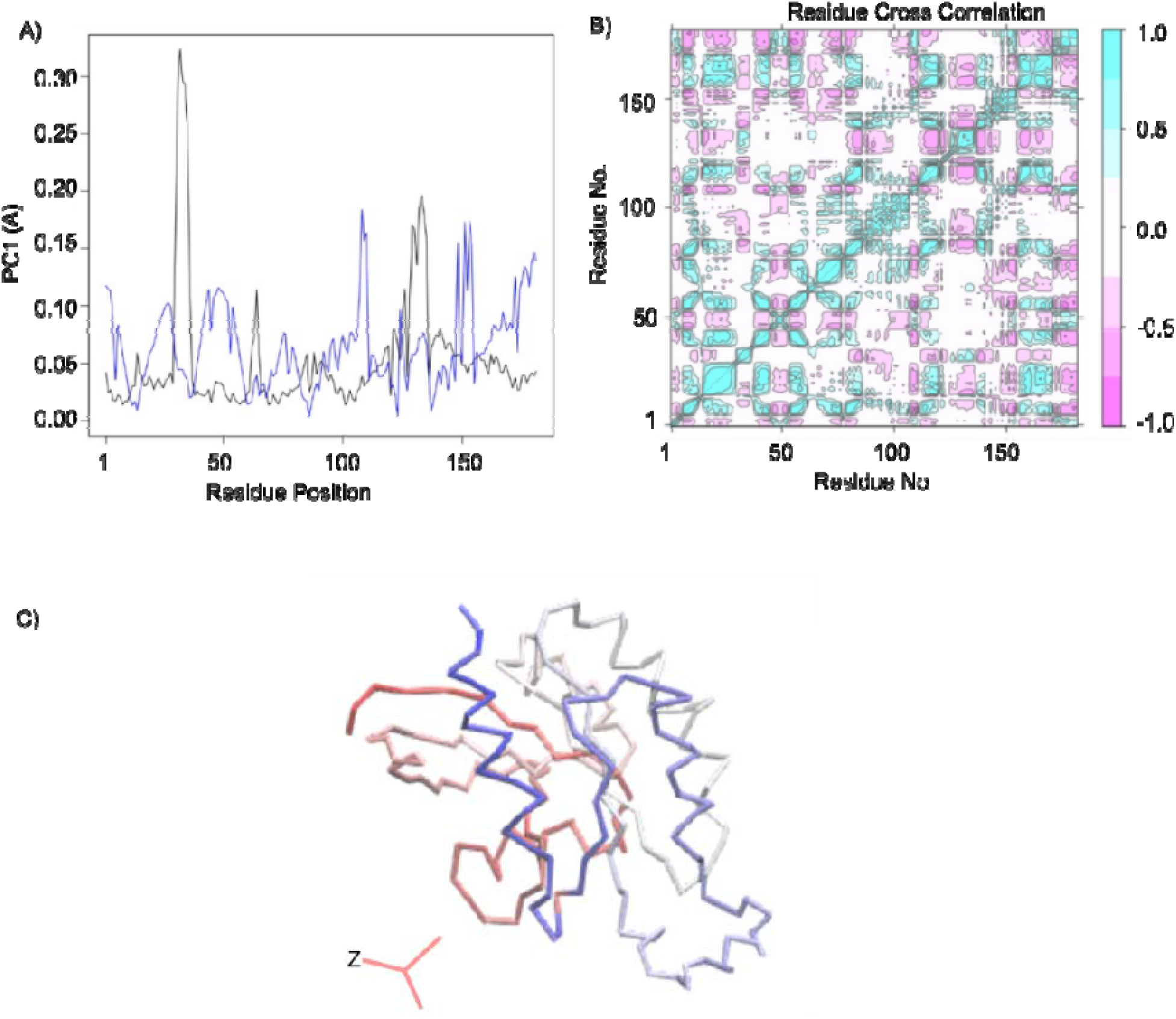
PC1 components and cross correlation analysis. **(A)** depicts an average fluctuation graph of all 181 components and the principal component. (**B**) Cross Correlation and Residue-wise loadings for PC1 (black) and PC2 (blue). Correlation analysis (Cij) of the motion during a 100ns MD simulation of the RhoB. (**C**) Contributions for first PC in a structural form and from the graph we acquired the knowledge of the contribution of pc1 was around 23% and one fourth in total.

Observing the scree plot of **Figure 8**, by ranking eigenvectors in order of their eigenvalues, highest to lowest, we got the principal components in order of significance [75]. The first PC accounts for more than one fourth; 23.34% of the overall variance. The second PC accounts for 17.95%. The first three components. together account for 50.88% ∼ 51%. From **Figure 8** we got picturesque information of three principal components against each other. The conformer plot of all selects is defined by PC1-PC3. Each point represents a structure and the point colour indicates the cluster id from a conformational clustering. Here, we can identify 4 coordinates and components clustered together. Highly correlated cells clustered together and such PCA plots are often used to find potential clusters [75]. The extent to which the atomic fluctuations/displacements of a system are correlated with one another can be assessed by examining the magnitude of all pairwise cross-correlation coefficients.

**Figure 8.**
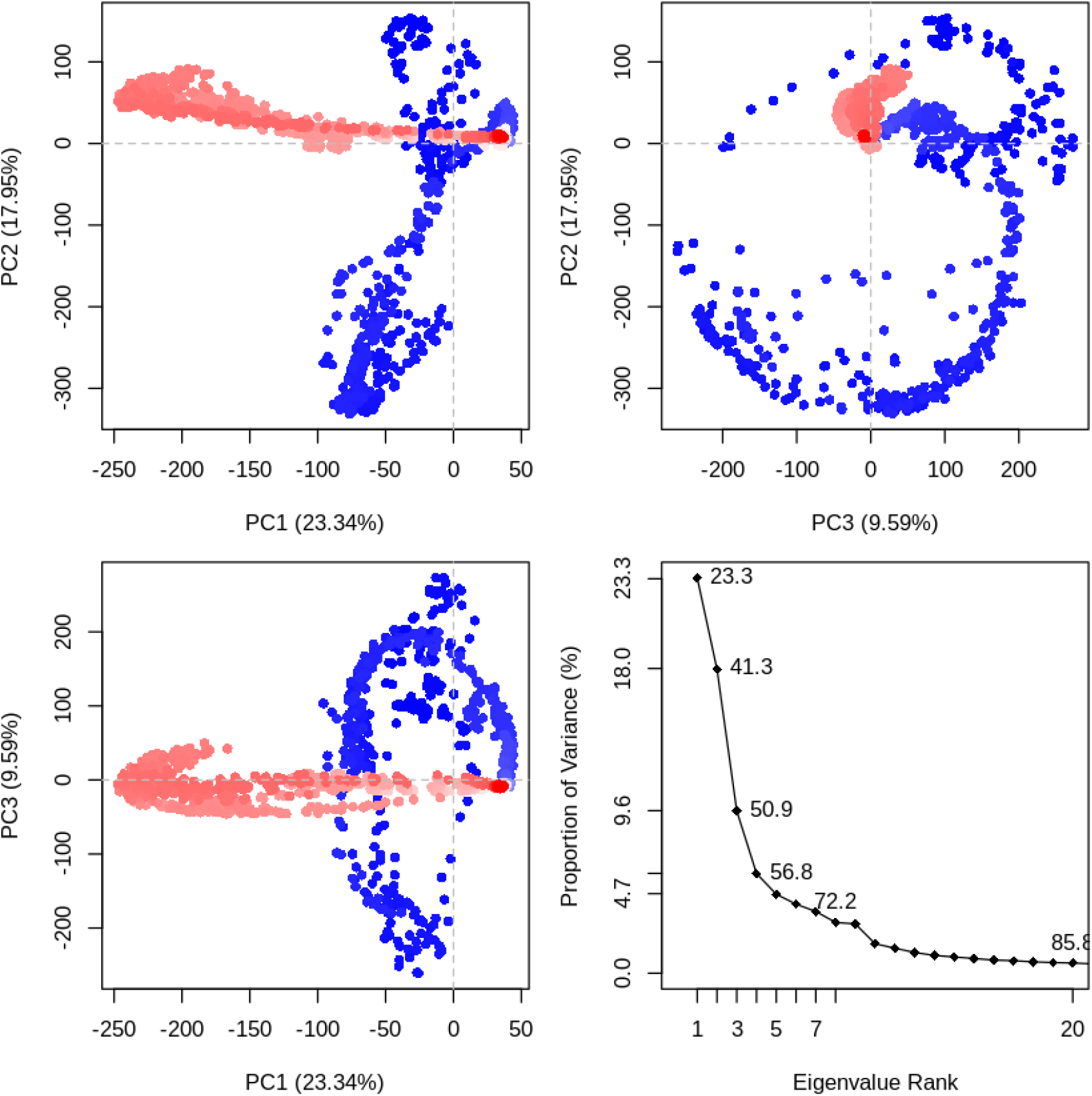
Principal Component Analysis. For the PCA, the last 100ns of each trajectory using the Bio3D package implemented in R. The first three eigenvectors, based on complex dominant motion, are extracted and compared. The variance captured by eigenvectors is also shown. Each point represents a structure and the point color indicates the cluster id from a conformational clustering. Projection of the trajectory onto the planes formed by the first three principal components. Conformers are colored according to the k-means clustering: PCA results for our Rho-B trajectory with instantaneous conformations (i.e. trajectory frames) colored from blue to red in order of time. The continuous color scale (from blue to whit to red) indicates that there are periodic jumps between these conformers throughout the trajectory. Below we perform a quick clustering in PC-space to further highlight these distinct conformers. In left side, PCA results for our Rho-B trajectory with instantaneous conformations (i.e. trajectory frames) colored from blue to red in order of time and in right side, Simple clustering in PC subspace.

## 4. Conclusion

In this investigation, we studied the physicochemical properties of Rho-related GTP-binding protein RhoB. Analysis of its primary sequence, which gave us cognition about its physicochemical characteristics, and functional activities. etc. It was acidic as its theoretical PI was 5.10 in the pH range, it was unstable because the value of the Instability index (II) was 46.35. Prediction of the secondary structure of this protein showed it was alpha helix dominating with 39.8% contribution in the construction of the secondary structure. The 3-D structure of a protein is biologically activated in nature since our query protein was uncharacterized along with no prior experimental structural information listed in PDB, we predicted its 3-D structure with a computational approach. An appropriate template 6hxu.1. A was selected among more than 30 templates from SWISS-MODEL based on its GMQE, QMAEN value with 100% identity, with 149 long amino acid sequences of the query protein on the other side Rho-related GTP-binding protein RhoB had 182 long amino acids in sequence. Validation of the model from the Ramachandran plot showed around 92% of amino acids occupied the most favoured region that indicating its reliability. Furthermore, to affirm the RhoB protein’s stability and compatibility, 100 nanoseconds (100,000ps) MD simulation was performed to know its atomic behaviour, conformational space, energies, RMSD, Radiation of gyration (Rg), RMSF with energies by using spc216 water model, OPLS AA force field and leap-frog integrator. Moreover, the RMSD value was reliable and can claim based on these values that the structure is suitable for advanced analysis. As we also analyzed and visualized the motion and movements of Principle components ignoring the rest reducing the noise and surface so that we can get over from complexity. These principal components can describe the characteristics of all the residues. Along with this we also calculated the SASA from webtool and trustable software expresses the accessibility of solvents of the protein residues. The hydrophobicity curve, residue x residue index, Cartesian coordinate of PC, and pairwise analysis of PC that was able to give use a better and valid understanding of the intramolecular characteristics of this protein residue. In conclusion, this investigation will assist the further studies involving the relation of the gene mutation and abnormalities induced by protein Rho-related GTP-binding protein RhoB in the progression apopotosis and so development of RhoB inhibitors. We performed a computational study on Rho-related GTP-binding protein RhoB and this study brought some limitations as well, for instance, homology modelled protein can’t reveal the exact structural features as the structure solved by the X-ray crystallographic method, Statistical knowledge comes before interpreting the extrapolated events of the protein and so further investigation is needed for the exact determination of physicochemical properties of this protein yet this computational study will be helpful for the future study of its clinical purpose.

## Acknowledgement

High-performance computing support for this research was provided by Science Outreach Servers.

